# Rapid life-history evolution reinforces competitive asymmetry between invasive and resident species

**DOI:** 10.1101/2023.10.25.563987

**Authors:** Elodie Chapuis, Philippe Jarne, Patrice David

**Affiliations:** MIVEGEC, Univ. Montpellier, CNRS, IRD, Montpellier, France; CEFE, CNRS − Université de Montpellier − IRD − EPHE, Montpellier, France

**Keywords:** invasion, competition, asymmetry, character displacement, laboratory, freshwater snails

## Abstract

Biological invasions by phylogenetically and ecologically similar competitors pose an evolutionary challenge to native species. Cases of character displacement following invasions suggest that they can respond to this challenge by shifting their traits. However, the actual impact of such shifts on competition are seldom studied. Here, we study competition between two freshwater snails from Guadeloupe (French Antilles), the native *Aplexa marmorata* and the introduced *Physa acuta*. The former has responded to invasion by rapid life-history evolution towards earlier maturity, higher fecundity and higher juvenile survival, traits that might favor rapid population growth in a noncompetitive context, but not necessarily in a competitive one. We here observe negative impacts of competition by both species on each other, though *P. acuta* is dominant and over generations largely displaces *A. marmorata* from co-cultures. In addition, our experiments suggest that *A. marmorata* populations having experienced competition by *P. acuta* for sufficient time in nature, have evolved to become even less tolerant to it. Though apparently paradoxical, this result supports the hypothesis that rapid life-history evolution has allowed *A. marmorata* not to resist competition, but to avoid it by increasing its specialization into the colonizer lifestyle previously documented by long-term field surveys. This example illustrates how evolution, in accordance with metacommunity coexistence theory, sometimes takes other ways than specialization into distinct types of resources or habitats to ensure coexistence between related species inhabiting the same landscape.

## Introduction

It is now commonplace to describe biological invasions as long-term, natural experiments (Sax et al., 2007; Davis, 2009) providing the opportunity to study the consequences of interspecific interactions on community dynamics and evolution. An invasive species develops a new network of interactions with residents, through for example predation, competition or habitat transformation (Carroll 2007; Davis, 2009; Prentis et al., 2008), thereby affecting their demography, abundance and selection regime (Simberloff 2013; Weber & Strauss, 2016). We here focus on competition, highlighting two important aspects with regard to species interactions. First, the strongest effects of competition are expected when the phenotypic and ecological similarity between invasive and resident species is high, for example when an invasive species meets with a phylogenetically related resident (the limiting similarity hypothesis; Duncan & Williams, 2002; Jeschke & Erhard, 2018). Second, biological invaders do not often wipe out the native species through competition, and most of the spectacular examples of extinctions following invasions are rather explained by predation or ecosystem engineering (Sax et al., 2007; Gallardo et al., 2016; David et al., 2017). Instead, introduced competitors will, most of the time, lead to a reduced abundance (Gallardo et al., 2016) and/or a narrowing of the niche of resident species (e.g., Duyck et al., 2006; Sidorovich et al., 2010). These two aspects make competition between related invasive and resident species a good situation to observe rapid evolutionary responses and their impact on community composition, combining a strong change in selection pressure and enough time to evolve. Phenotypic shifts due to microevolution and/or plasticity may then contribute to modify the intensity of reciprocal competitive impacts between invasive and resident species, as shown in plants (Huang et al., 2018; Germain et al., 2020). However most experimental studies generally look at trait change and do not directly measure competition (Stuart et al., 2014; Alexander et al., 2015; Hart et al., 2019). Here we study under laboratory conditions the competition between two related species, one invasive and one resident, and how previously documented evolutionary changes in traits during invasion (Chapuis et al., 2017) modify this competition.

At least two general predictions have been proposed on the impacts of competitive interactions between invasive and resident species, based on the idea that coevolution of competing species contributes to long-term coexistence (Lankau, 2009; 2011; Faillace & Morin, 2016). A first prediction is the existence of an initial competitive asymmetry when two generalists, related, invasive and resident species first come into contact. The species have not had the opportunity to coevolve and mitigate their reciprocal impacts (for example, through some form of niche specialization or character displacement; Freckleton & Watkinson, 2001), so that one of the two may often largely dominate the other on the most abundant resource, an interaction more asymmetrical than would be expected in a pair of long-term sympatric competitors. However, among all possible candidate invaders, those that are competitively superior to the resident will invade with a greater probability (the ‘invasion filter’: David et al., 2017; Graiger et al., 2019). As a result, we might expect that successful invaders will often be largely dominant over related residents, possibly restricting them to marginal resources or niches that they are unable to exploit themselves (Lagos et al., 2017). The second prediction, which is a consequence of this initial asymmetry, is that the strongest selection pressure is expected to bear mostly (though not exclusively; see Jones & Gomulkiewicz, 2012) on the resident species. Two evolutionary trajectories can then be envisaged: (i) the resident species evolves to resist competition, becoming more competitive on the resource occupied by the invasive species (Germain et al., 2020); (ii) it evolves to avoid competition, through ressource shift and character displacement (Pfennig & Pfennig, 2009). These two possible evolutionary trajectories (resistance and avoidance) can also be related to the two broad mechanisms, equalizing and stabilizing, promoting species coexistence in Chesson’s theory (Chesson, 2000). Equalizing mechanisms tend to reduce average fitness differences between species within their common environment, while stabilizing mechanisms are present when intraspecific competition exceeds interspecific competition, e.g., because species specialize on different resource ranges, resulting in negative frequency-dependence and preventing competitive exclusion. Interestingly, resisting and avoiding competition may require different sets of traits in trade-off with each other, or different values of the same trait, suggesting disruptive selection towards traits promoting either tolerance, or avoidance. The direction taken by the evolutionary response partly depends on the initial asymmetry (Jones & Gomulkiewicz, 2012). Under a very asymmetrical interaction (invasive very dominant), the probability of resistance is low and the most likely response will be avoidance, while a small initial asymmetry would increase the probability to evolve resistance (Huang et al., 2018; Germain et al., 2020).

Phenotypic evolution in pairs of competing species has classically been studied through the concept of character displacement (Tobias et al., 2014). Such studies mostly focus on traits linked to the classical niche concept, i.e., traits involved in the exploitation of different food resources (e.g., beak morphology in Darwin’s finches; Grant & Grant, 2006) or different microhabitats (e.g., limb structure in *Anolis* lizards; Feiner et al., 2021). A relatively under-explored category of traits, though potentially very important in this context, is life-history traits. These traits may be subject to rapid evolution after contact between invasive and resident species both because they often have a high genetic variance and evolvability relative to morphological traits (Houle 1992), and because modern coexistence theory has highlighted their potential to promote coexistence on the same type of resource, through metapopulation mechanisms, such as colonization-competition trade-off (Higgins & Cain, 2002) or competition-persistence trade-off (Muller-Landau, 2010). Accordingly, some life histories can favor persistence when the resource is low and stable, while others favor colonisation and population growth in patches of abundant and ephemeral resources. For example, in plants, the life-history trade-off theory has been used to study traits linked to seed dispersal (Turnbull et al., 2004; Cadotte et al., 2006). In the context of coexistence theory based on life-history trade-offs, the alternative between resistance to competition versus avoidance of competition translates into convergence versus divergence of life-history profiles along the colonization-competition gradient.

Here we test these predictions (resistance/avoidance) by focusing on a pair of generalists, related species (freshwater snails), one invasive and the other resident. We previously observed that populations of the resident species have recently evolved towards a life-history profile that seems less competition- and more colonization-oriented, after contact with the invasive (Chapuis et al., 2017). We measure the competitive interactions between these species in the laboratory to test (i) whether the invasive species is indeed competitively dominant (competitive asymmetry), (ii) whether the evolution of life-history traits in the resident has modified competitive impacts and in which direction, and (iii) to assess whether the competitive interaction has evolved in a population-specific way. These three hypotheses are summarized in Table 1.

**Table 1.** Main hypotheses and associated predictions in our experiments. They are discussed in Introduction, and how the experimental tests were conducted in Material and methods. Note that Front and Core populations of the invasive species *P. acuta* showed relatively little trait differences, except for a slight decrease in fecundity (Chapuis et al. 2017) in the Front populations, assumed to reflect temporary founder effects. We therefore do not have clear expectations about differences in the outcomes of competition experiments – Front populations might, if anything, be slighlty less harmful competitors than Core ones to the resident species. We nevertheless kept the Front / Core distinction in the statistical analysis to check for a possible effect.

| Hypotheses | Predictions |
| --- | --- |
| (i) Initial competitive asymmetry favouring the introduced species => *P. acuta* dominates. | (a) The impact of *P. acuta* on traits associated with fitness of *A. marmorata* is more negative than the reverse. Individuals of both species have lower growth and/or survival when their competitors are *P. acuta* than when these competitors are replaced by the same number of *A. marmorata* individuals.<br>(b) The relative recruitment of *P. acuta* in mixed populations is higher than that of *A. marmorata*, so that the frequency of *P. acuta* increases over the generations. |
| (ii) Trait evolution in the resident species *A. marmorata* reflects an avoidance of competition rather than an increased tolerance to competition. | Growth and/or survival of Experienced populations (E) of *A. marmorata* are more negatively affected by *P. acuta* than those of Naive populations (N): Experienced are less tolerant to competition than Naive. |
| (iii) *A. marmorata* and *P. acuta* have locally coevolved. | (a) If coevolution reinforces competitive impacts: survival/growth in one species are more negatively affected by sympatric competitors of the other species than by allopatric competitors.<br>(b) If coevolution mitigates competitive impacts: traits are more negatively affected by allopatric competitors than by sympatric ones. |

We take advantage of the spatial structure of the invaded region, with sites ‘anciently’ versus ‘recently or not yet invaded’, at the time of sampling – the latter acting as evolutionary controls. However, spatial structure potentially adds another source of complexity: competition is by definition a local interaction and, if gene flow is too slow to homogenize populations at the time scale considered here (a few generations or tens of generations), evolution may take place, to some extent, independently in different invaded populations. In this situation, competitive interactions might differ depending on whether we confront invasive and resident populations from the same site versus from different sites.

We conducted laboratory experiments with a pair of pulmonate freshwater snails of the same family (*Physidae*). In Guadeloupe (Lesser Antilles) the native species (*Aplexa marmorata*) occupies a network of freshwater ponds that have been recently invaded by the ecologically and phylogenetically (Wethington & Lydeard, 2007) related species *Physa acuta*, a cosmopolitan invasive species (Bousset et al., 2004; Ebbs et al., 2018). Owing to a long-term yearly metacommunity survey we have a very detailed knowledge of invasion of new sites by *P. acuta* since 2001, allowing us to contrast long-invaded versus not or recently invaded sites. Metapopulation models on presence/absence data (Dubart et al., 2019; Pantel et al., 2022), time-series of changes in species abundance within sites (Chapuis et al., 2017) and joint species-distribution models at the metacommunity scale (Dubart et al., 2019, 2022) all detected negative and reciprocal impacts of competition in the field between *P. acuta* and *A. marmorata* during the 2001-2017 period. No loss of neutral genetic variability was detected at the invasion front in *P. acuta*, but it decreases in *A. marmorata* populations that have been confronted by *P. acuta* for a longer time (Jarne et al., 2021). In addition, as mentioned previously, a common-garden experiment showed that life-history traits in *A. marmorata* evolve in a few years (less than two) after a population is reached by *P. acuta*, becoming mature sexually earlier and more fecund (Chapuis et al., 2017). We conducted laboratory competition experiments with the same set of populations than Chapuis et al. (2017).

## Material and methods

### Biological model

Our field sites are freshwater ponds or intermittent creeks located in Grande-Terre and Marie-Galante (Guadeloupe; see Figure 1). The native species *A. marmorata*, and the invasive *P. acuta* belong to a community of around 25 species of freshwater snails, all detritivorous and herbivorous, that collectively constitute the most important biomass of the macrobenthos in these habitats (Pointier, 2008; Pantel et al., 2022, Dubart et al., 2022). *Physa acuta* was first detected in Guadeloupe in the late 1970s and has recently experienced a spectacular spread from two sites occupied in 2001, to more than 80% of the ca. 280 surveyed sites in 2020. The first report of *P. acuta* in Marie-Galante is more recent (2010). The native *A. marmorata* also occupies a large fraction of these sites with a moderate temporal decrease in this occupancy, but a significant decrease in population density, coinciding with the arrival of *P. acuta* (Chapuis et al. 2017). This hermaphroditic species self-fertilizes at rates exceeding 90% in natural populations (Dubois et al., 2008; Escobar et al., 2011), while *P. acuta* is mainly outcrossing. In both species sexual maturity of isolated individuals is reached at about four to five weeks under laboratory conditions (24°C, 12 hours of light; Chapuis et al., 2017). Egg capsules include up to 30 eggs inserted in a gelatinous matrix, and are laid on hard surfaces (e.g., branches or stones in natural populations, box walls in the laboratory). Juveniles hatch after about a week. Growth is continuous, and adults reach a size of about 10-15 mm in shell length. The generation time under natural conditions is estimated to be around two months, meaning 5-6 generations per year. Individuals can live up to six months under laboratory conditions.

**Figure 1.**
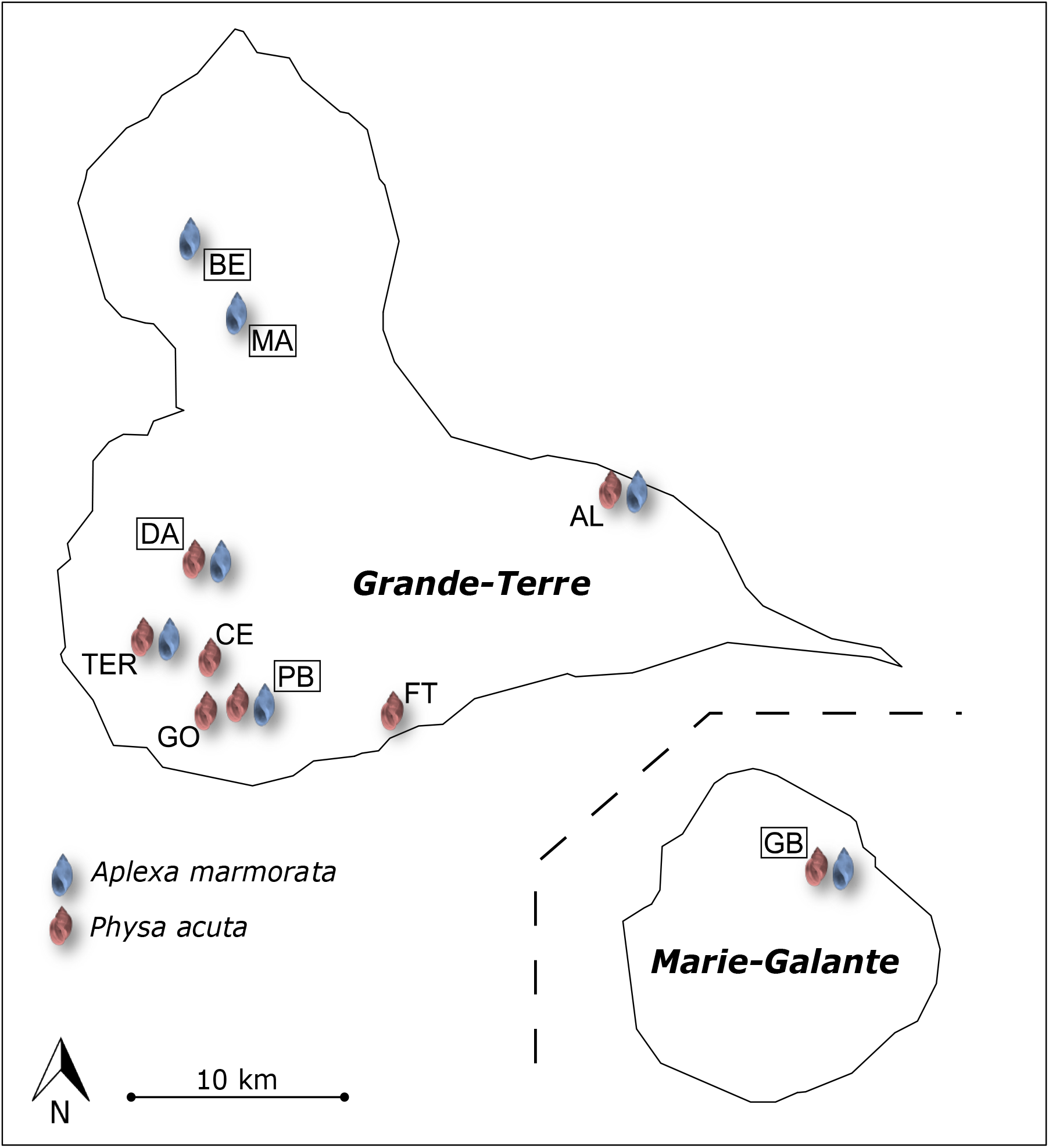
Distribution of the 10 sites sampled in Guadeloupe for our experiment. Snail populations are represented by colored pictograms: blue for *Aplexa marmorata* and red for *Physa acuta*. N (never or newly-invaded) populations of *A. marmorata* are indicated by a box. See Table 1 for the acronyms of population names. The islands of Grande-Terre and Marie-Galante, which appear exaggeratedly close here, are in reality around 30 km apart.

### Field collection

We sampled G_0_ snails from 10 sites: TER, AL, GB, DA, CE, MA, FT, PB, G0, BE (Figure 1, Table 2 and Chapuis et al., 2017 for more details on populations). Seven sites harboured populations of *A. marmorata* and eight *P. acuta*, and five were occupied by both species. Sampling was done in January 2010. Individuals were brought back to the laboratory in Montpellier (France) and the experiment was carried out on the second laboratory generation in constant conditions (12/12 photoperiod and 25°C). Populations of *A. marmorata* were classified into two types depending on when/whether *P. acuta* invaded the site, based on previous analyses (Chapuis et al., 2017): ‘naive’ (N) refers to populations where, at the time of sampling, *P. acuta* had never been observed or had been observed only recently (less than two years) and ‘experienced’ (E) to populations that have evolved in presence of interspecific competition as the first observation of *P. acuta* had taken place more than two years before sampling (Table 2). On the other hand, populations of *P. acuta* can be classified as ‘front’ (invasion front, F) for populations present for less than two years in a site and ‘core’ (C) when present for more than two years (Table 2). Note that the contrast between F and C populations in *P. acuta* is of a different nature from that between N and E populations. In *P. acuta*, F populations are younger (founded more recently) than C populations, while N and E populations in *A. marmorata* are all anciently established. In addition, F populations are not ‘naive’ as their ancestors have been exposed to the same history of interspecific competition as C populations (they were recently founded, but the founders were coming from C populations). An additional distinction refers to the fact that competing populations in mixed cultures could have been sampled in the same site (hereafter ‘sympatric’) or in different sites (‘allopatric’) - we referred to this effect as origin of the competitor.

**Table 2.** The 10 sites in which *A. marmorata* (native) and *P. acuta* (invasive) were sampled. The year during which *P. acuta* was first detected (as of 2010 sampling campaign) is given in the ‘first detection’ column (‘never’ indicates that *P. acuta* was never detected). The invasion status is given for each population (N for naive, E for experienced in *A. marmorata*; F for front and C for core in *P. acuta*, details in text). Hyphens (--) in the last two columns indicate that the species was not sampled in this site.

| Site | Acronym | First detection of *P. acuta* | *A. marmorata* | *P. acuta* |
| --- | --- | --- | --- | --- |
| Belin | BE | Never | N | -- |
| Malakoff | MA | Never | N | -- |
| David | DA | 2010 | N | F |
| Grand-Bassin | GB | 2010 | N | F |
| Port Blanc | PB | 2010 | N | F |
| Fond Thezan | FT | 2007 | -- | C |
| Alleaume Nord | AL | 2004 | E | C |
| Caraque Est | CE | 2004 | -- | C |
| Goudenave | GO | 2004 | -- | C |
| Terrasson | TER | 2004 | E | C |

### Laboratory experiments

We collected individuals (G_0_) from each population in January-February 2010, and they were brought back to the laboratory in Montpellier to lay eggs (G_1_). These G_1_ individuals were used in a previous experiment (Chapuis et al., 2017) and their offspring (G_2_) were used for the following experiment. The G_2_ individuals were raised separately in 80 mL plastic boxes from hatching to three weeks (Chapuis et al., 2017). The competition experiment took place in June-August 2010 as follows: five three-week old G_2_ individuals from a given population of each species were put together in a 1L tank (i.e., 10 individuals per tank). Once mixed, snails of the same species could no longer be identified individually, but interspecific differences were sufficient to distinguish adult individuals from each species. Among all possible pairs of populations, we were able to constitute 15 pairs (see Table 3). We chose to set up both allopatric and sympatric pairs when possible (sympatry is when the two populations originated from the same site, and allopatry when they originated from different sites). It should be noted that, by definition, it is not possible to establish sympatric pairs for *A. marmorata* populations that have never been invaded by *P. acuta*. Among the allopatric pairs, we could make E/C, N/C, and N/F combinations. However, the low number of *A. marmorata* E populations and of *P. acuta* F populations did not allow us to make the E/F combination. For each population and each species, control tanks were set up and included 10 individuals from only one species and population. Two replicates were made for each type of tank, giving a total of 60 experimental tanks (Table 3).

**Table 3.** Combinations of populations (for acronyms see Table 2) used in the interspecific competition experiment. The 15 pairs (indicated by A or S) included seven *Aplexa marmorata* populations and eight *Physa acuta* populations. A = allopatric pair and S = sympatric pair; N for naive, E for experienced in *A. marmorata*; F for front and C for core in *P. acuta* as in Table 2. Cells are shaded for the pairs that were not constituted.

| *Physa acuta* | AL<br>(C) | TER<br>(C) | CE<br>(C) | FT<br>(C) | GO<br>(C) | GB<br>(F) | PB<br>(F) | DA<br>(F) |
| --- | --- | --- | --- | --- | --- | --- | --- | --- |
| *Aplexa marmorata* |  |  |  |  |  |  |  |  |
| AL (E) | S | A |  |  |  |  |  |  |
| TER (E) | A | S |  |  | A |  |  |  |
| GB (N) |  |  |  |  |  | S | A |  |
| PB (N) |  |  |  |  |  | A | S | A |
| DA (N) |  |  |  |  |  |  |  | S |
| BE (N) |  |  | A | A |  |  |  |  |
| MA (N) |  |  | A | A |  |  |  |  |

#### First stage of the competition experiment

The experiment was initiated using three-week old individuals (i.e., before sexual maturity which occurs around 4-5 weeks under our laboratory conditions; Chapuis et al., 2017), and consisted of two successive stages. The first stage (from grouping to 35 days / five full weeks later) mostly focused on the effects of competition on adult life-history traits. G_2_ individuals of both species interacted freely for five weeks and were then discarded. In snails, growth and other traits depend on population density (Dan and Railey, 1982; Dillon, 2000) so the same initial density of 10 individuals per tank was used for all tanks including intraspecific controls. However, once the experiment started, we chose not to replace dead individuals to mimic “natural situations”. Shell length was measured two and five weeks after the initial grouping. The number of surviving G_2_ adults of each species was counted at week 5. At that time, G_2_ adults had started to reproduce and were living together with their progeny (egg masses or hatchlings). Adults tend to monopolize resources and inhibit the growth of juveniles, and were then removed at week 5.

#### Second stage of the competition experiment

In the second stage (weeks 5 to 16), we focused on relative recruitment of the two species under interspecific competition. To this end, we kept only aquaria where the two species had been initially introduced. Juveniles of both species developed under intense competition, both within and among species. Competition went on for 11 weeks, after which we measured the number of G_3_ snails recruited for each species within each aquarium (week 16). Before they reach a certain size (∼4 mm), snails cannot easily be identified to the species level. We therefore decided to consider only individuals over this threshold size as recruited in the next generation. Note that some of the recruited G_3_ snails may have already started to reproduce at week 16. The relative recruitment of *P. acuta* and *A. marmorata* within each aquarium represents an integrative outcome of competition over two generations (i.e., relative population growth rate, accounting for interspecific differences in fecundity, juvenile survival, and growth).

Experiments in the two stages were conducted as in previous work (e.g., Escobar et al. 2011, Noel et al. 2016). Snails were kept in 12L:12D photoperiod in a dedicated room (‘molluscarium’) at a 24±1ºC temperature. Once a week, water was changed and snails were fed with boiled and ground lettuce. In aquaria, the lettuce was provided twice a week in moderate quantity, so that it was completely consumed before the next feeding, thus enhancing competition.

### Data analysis

Our data <u>(10.5281/zenodo.10041279</u>.) were used to test our main predictions (see Introduction and Table 1) based on statistical analyses conducted separately for each species (<u>10.5281/zenodo.10041344</u>.). The software R and the “lme4” package in R (Bates et al., 2015) were used for all statistical analyses (R Core Team, 2019).

Shell size (week 2 and 5) was analysed in two steps. In the first step, we focused on competition asymmetry (difference in size under intraspecific versus interspecific competition), and its potential dependency on the status of the population (N versus E for *Aplexa marmorata*; F versus C for *Physa acuta*).

We fitted linear mixed models with fixed effects “type of competition” (intraspecific in control tanks, interspecific in mixed tanks), population status, and their interaction. Population (in interaction with type of competition) and replicate (nested within population-by-competition combination) were included as random effects. Model simplification and likelihood-ratio tests were used to test the significance of fixed effects (keeping random effects unchanged). When the interaction was significant the effect of type of competition was tested within each status separately. In the second step, we kept only mixed tanks (i.e., interspecific competition) and focused on possible differences in competitive impact depending on coevolutionary history (allopatric versus sympatric pairs of populations, i.e., sampling site effect). We fitted linear models with a single fixed sampling site effect; random effects were again population (in interaction with sampling site) and replicate. The same two-step analysis was performed on survival at week 5, using a generalized linear mixed model (GLMM) with binomial error, with the random ‘replicate’ effect accounting for overdispersion (i.e., excess of residual variance among aquaria).

The recruitment of juveniles of the two species (after 16 weeks) was measured only in mixed aquaria. First, we analysed, for each species separately, the number of juveniles recruited using GLMMs with Poisson error, with status (E/N or F/C depending on species), sampling site and their interaction as fixed factors, and population and replicate as random factors (the replicate factor accounts for overdispersion, as previously). Second, we focused on relative recruitment of juveniles of the two species using binomial GLMMs on the numbers of recruits per species. The fixed factors included were the same as previously, while the random factors were population pair and replicate.

Size, survival and recruitment were used to test the predictions in Table 1, in which we indicated the expectations and tests on these traits.

## Results

### Body size

#### Physa acuta

In *P. acuta*, we found a (marginally) significant status effect on shell length at both week 2 and week 5, as individuals from F populations were slightly larger than those from C populations (Table 4; Figure 2). Individuals exposed to interspecific competition were, on average, similar in size to those exposed to intraspecific competition (Table 4; Figure 2). However, the outcome of interspecific competition depended on the origin of *A. marmorata* competitors (Table 4): snails exposed to sympatric competitors grew smaller than those exposed to allopatric competitors (Figure 3).

**Table 4.**
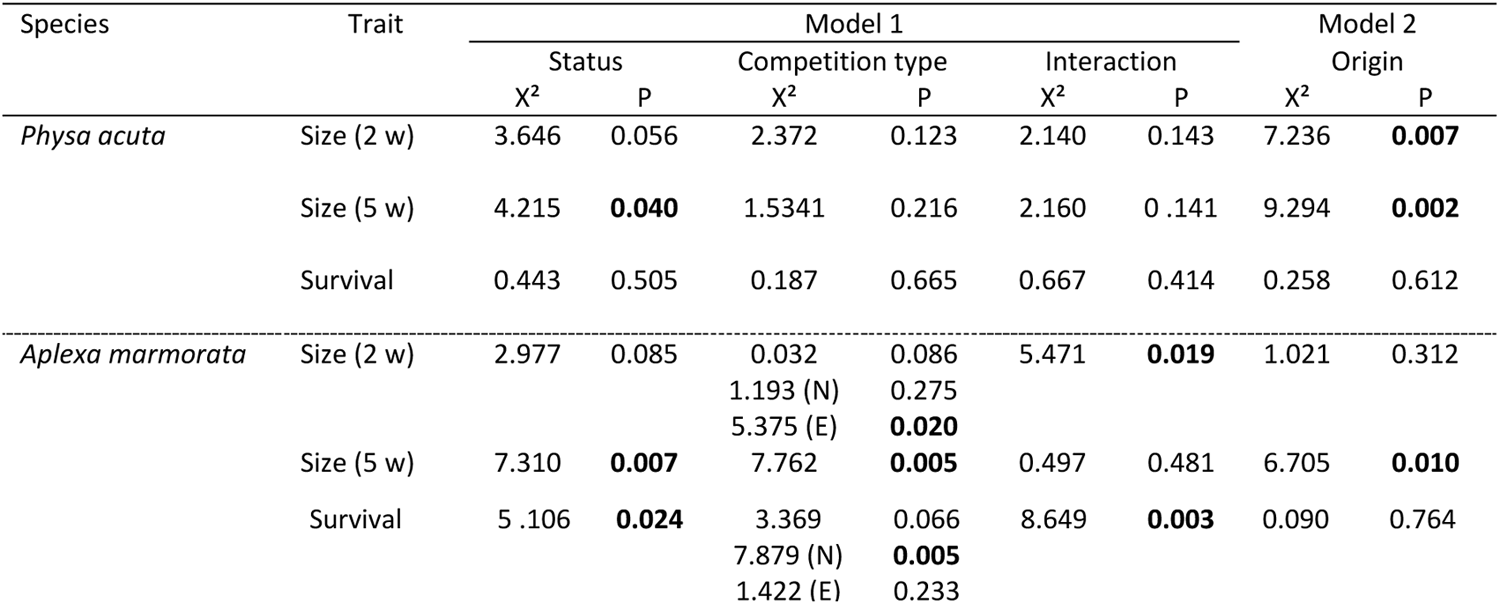
Statistical tests on size and survival of G_2_ individuals in the two species studied. Size was estimated at week 2 and 5, and survival between weeks 0 and 5. In model 1, we considered all experimental units (one- and two-species cultures), their status (core and front in *P. acuta*, and naive and experienced in *A. marmorata*), the competition type (intra- and interspecific) and their interaction. N and E refer to tests of competition type made separately within naïve or within experienced populations, performed when the interaction was significant. Only two-species cultures were considered in Model 2, in which we tested the effect of the origin of the competitor (sympatric versus allopatric). All linear and generalized linear mixed models included population and replicate as random factors. Size was modelled with a Gaussian distribution, and survival with a Binomial one. We report the value of the chi-square test (X^2^) with the associated probability value (P). All tests with one degree of freedom. In bold, P<0.05.

**Figure 2.**
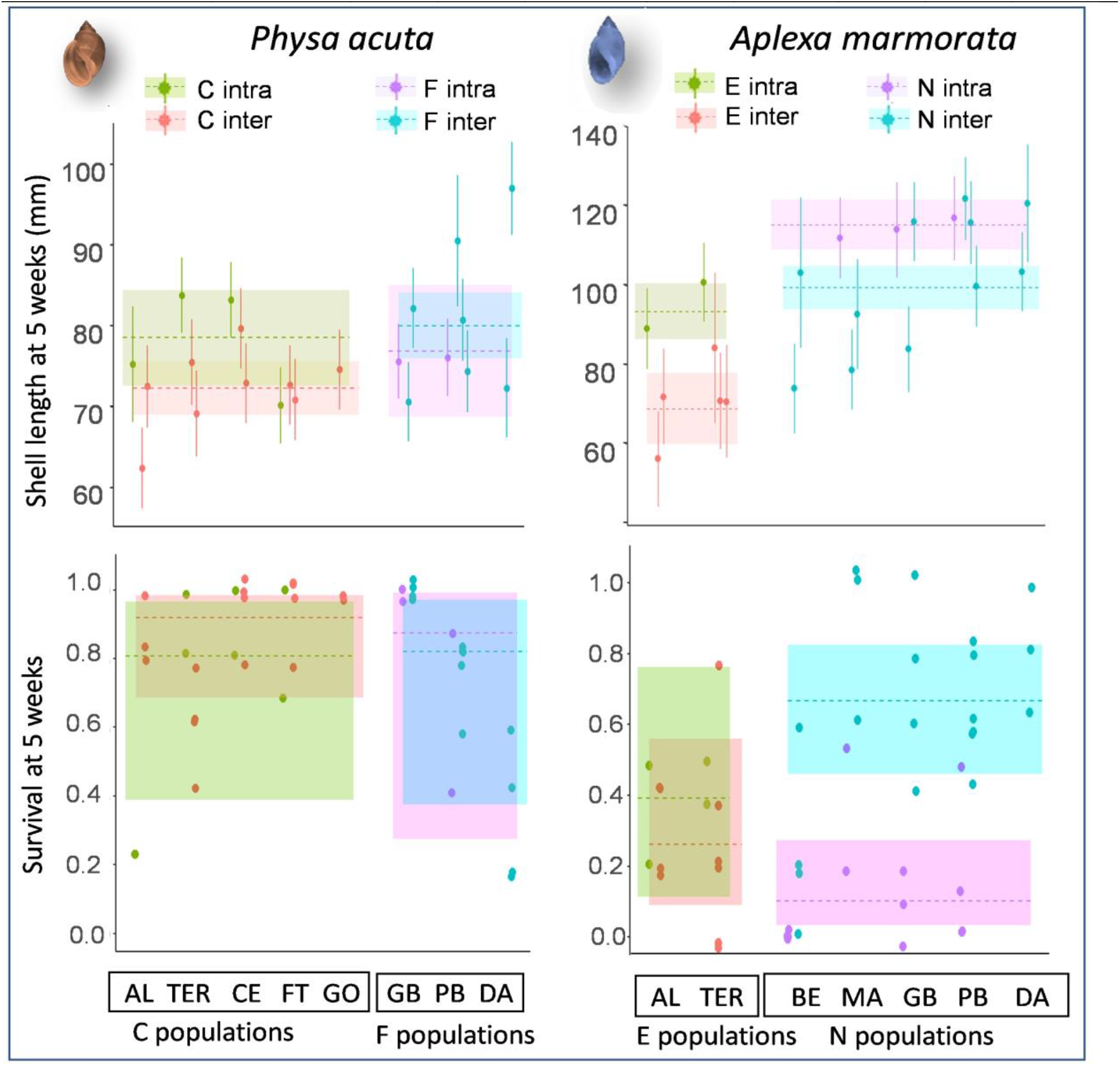
Effect of population status and competition type (intra-versus interspecific) on G_2_ shell size and survival (at week 5) in the resident species *Aplexa marmorata* (right) and in the invasive one *Physa acuta* (left). This figure corresponds to Model 1 in Table 4. Populations are indicated at the bottom (arranged by status: Front - F, Core - C for *P. acuta*; Naive - N, Experienced - E for *A. marmorata*), with full names in Table 2. Each point for shell length represents the mean of the focal population of the focal species when competing with itself (intraspecific competition, green or purple) or with a population of the other species (interspecific competition, red or blue), with error bars (+/- 1 standard error on the mean) based on both among-individual and among-replicate variance. Survival is represented by dots for each replicate (one replicate is 10 individuals; survival is therefore a multiple of 0.1 but dots have been slightly jittered to facilitate visualisation). Horizontal dotted lines indicate model estimates for each status and competition type, with shaded zones indicating 95% confidence intervals. Note the difference in scale between the two species, for shell size. Refer to Table 2 for combinations of population pairs in interspecific interactions.

**Figure 3.**
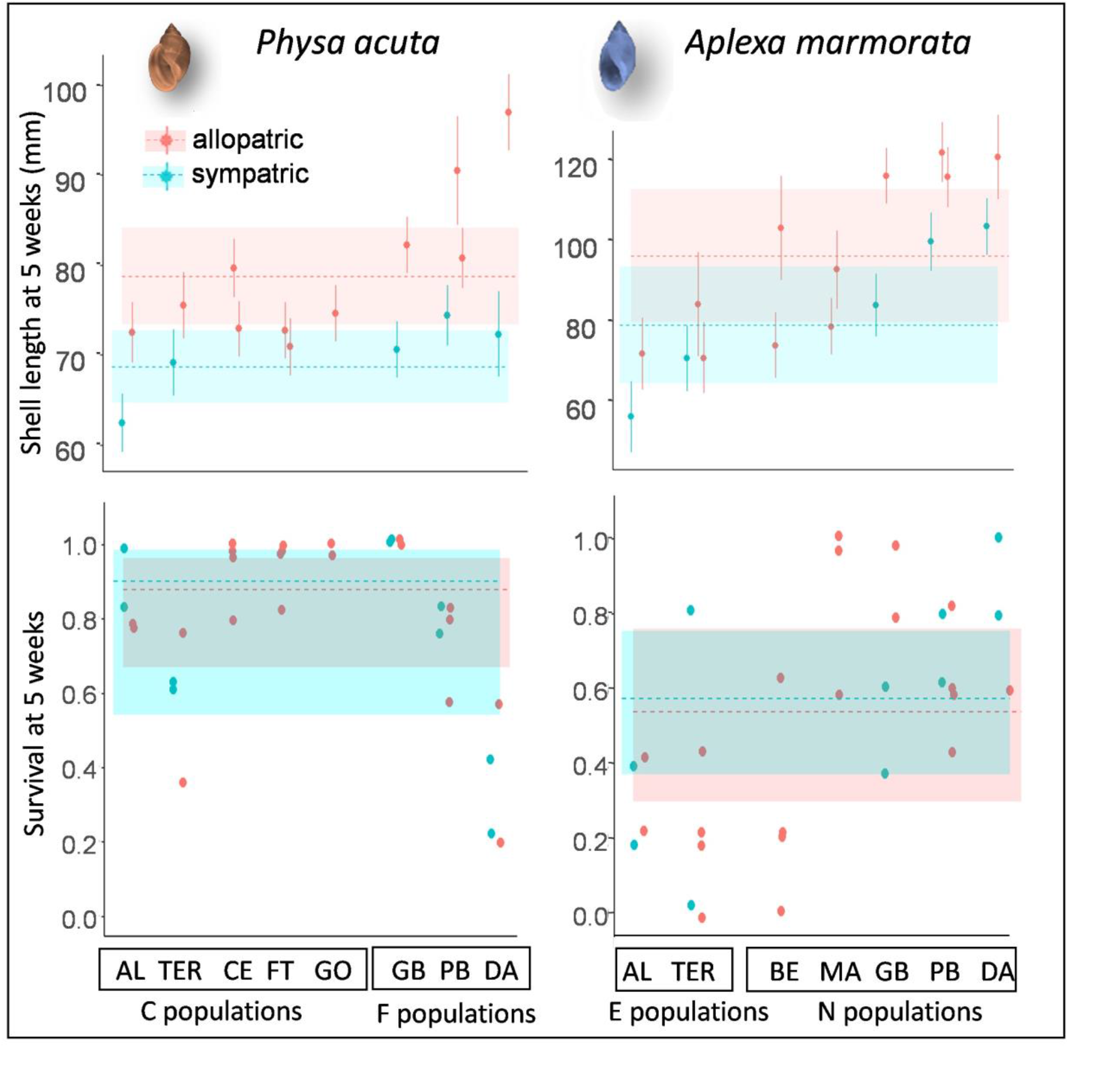
Effect of site origin (sympatric versus allopatric) on G_2_ shell size (at week 5) and survival of the two snail species, the invasive one *Physa acuta* (left) and the resident one *Aplexa marmorata* (right), under interspecific competition. This figure corresponds to Model 2 in Table 4. Populations and replicates are the same, and depicted in the same way, as in figure 2. However, only situations of interspecific competition are represented, and colors (blue versus red) indicate the sympatric or allopatric origin of the competitors (whether they come from the same site as the focal population or not). Horizontal dotted lines indicate model estimates for each origin, and shaded areas 95% confidence intervals (we used the model without interaction between competitor origin and population status as this interaction was never significant, so a single estimate for each competitor origin is represented). Note the difference in scale between the two species, for shell size.

#### Aplexa marmorata

At 5 weeks, shell size was significantly affected by both population status and the type of competition in *Aplexa marmorata* (Table 4; Figure 2): snails from N populations were bigger than snails from E populations, and snails grown under interspecific competition were smaller than snails grown in single-species cultures. The effect of interspecific competition was more pronounced, resulting in smaller snails, when competitors were sympatric than when they were allopatric (Table 4; Figure 3). As for *P. acuta*, this difference was observed in each of the five populations where the comparison could be made. The same tendencies were already observed at week 2, with slight differences. First, the effect of competition type was significant only in E populations, where interspecific competition reduced size relative to intraspecific competition, resulting in a significant competition-by-status interaction. Second, the effect of allopatry versus sympatry was not significant (Table 4).

### Survival and recruitment

#### Physa acuta

In *P. acuta*, the survival of the adult G_2_ individuals was not significantly affected by population status, type of competition, or the allopatric versus sympatric origin of competitors (Table 4; Figures 2 and 3). The number of G_3_ recruits of *P. acuta* at the end of the experiment in mixed cultures was not significantly affected by the origin of competitors, but the status effect was marginally significant (Table 5), as C populations recruited on average more offspring than F populations (although recruitment was highly variable in both cases; Figure 4).

**Table 5.** Effect of population status (detailed in text and in Table 2), origin of the competitor (allopatric versus sympatric) and interaction of both on the recruitment of G_3_ individuals at week 16 in two-species cultures. Population and replicate were included as random factors. Models were Poisson-distributed GLMMs, and we report the value of the chi-square test (X^2^) with the associated probability value (P). All tests with one degree of freedom. In bold, P<0.05.

| Species | Status |  | Origin |  | Interaction |  |
| --- | --- | --- | --- | --- | --- | --- |
|  | X<sup>2</sup> | P | X<sup>2</sup> | P | X<sup>2</sup> | P |
| *Physa acuta* | 4.375 | **0.036** | 1.498 | 0.221 | 1.545 | 0.214 |
| *Aplexa marmorata* | 0.058 | 0.809 | 0.003 | 0.954 | 1.498 | 0.221 |

**Figure 4.**
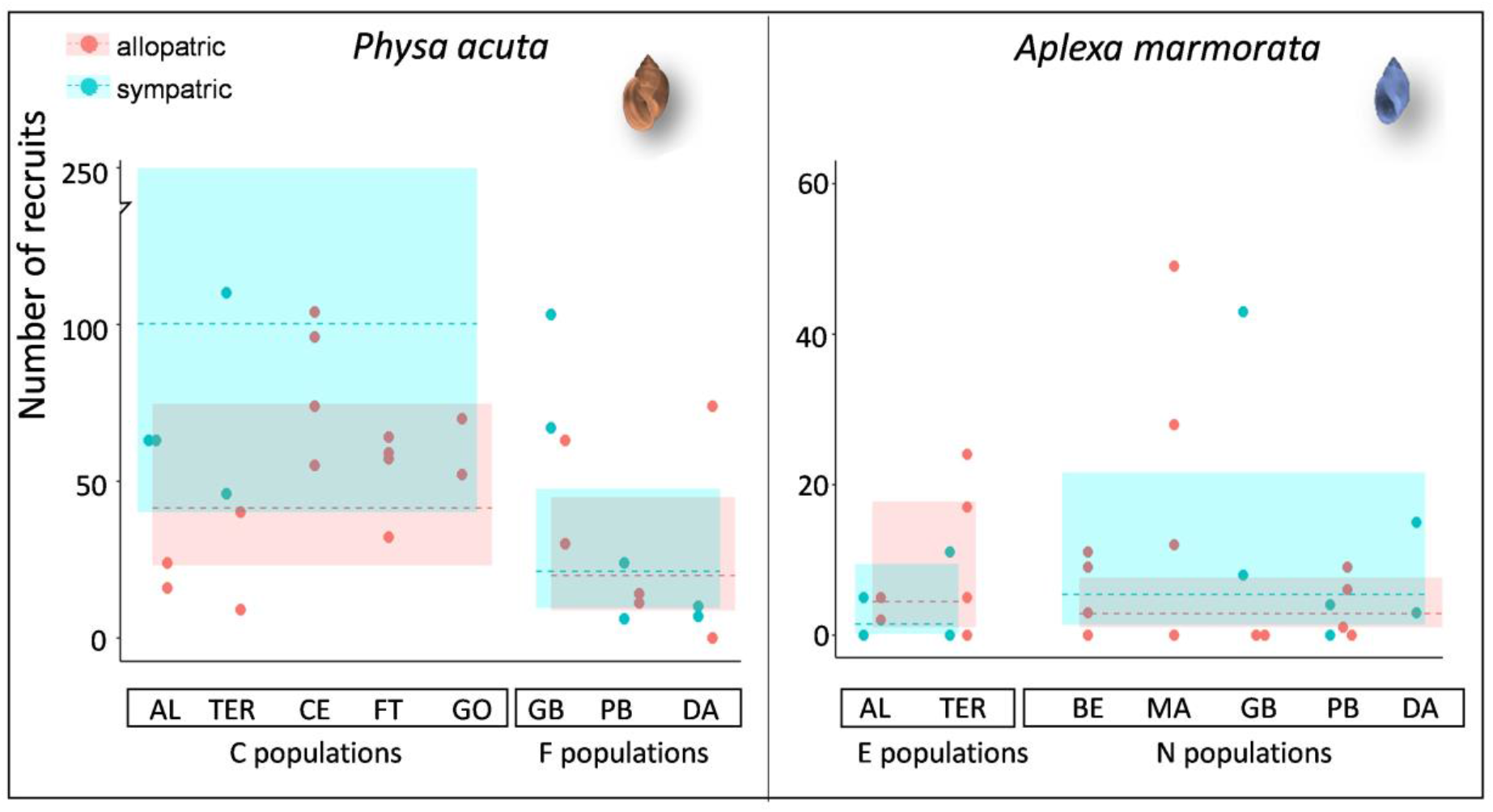
Effect of population status and sampling site (sympatric versus allopatric) on recruitment (at week 16) of the two snail species, the invasive one *Physa acuta* (left) and the resident one *Aplexa marmorata* (right), under interspecific competition. Populations are indicated at the bottom of each graph (arranged by status: Front - F, Core - C for *P. acuta*; Naive - N, Experienced - E for *A. marmorata*), and full names in Table 2. Each dot represents a replicate. Horizontal dotted lines indicate model estimates (Poisson GLMM) for each status and competition type, and shaded areas denote the corresponding 95% confidence intervals. Note the difference in scale between the two species. The data represented here are a subset of those in Figure 2 (interspecific interactions only).

#### Aplexa marmorata

In *A. marmorata* the survival of G_2_ individuals at week 5 was affected by the type of competition, but the effect differed depending on population status (significant interaction in Table 4). In E populations, *A. marmorata* survived at similar rates whether in intra or interspecific competition, while in N populations, they survived better under interspecific competition than in pure cultures (Table 4; Figure 2). Under interspecific competition, there was no difference in survival depending on whether the competitors were allopatric or sympatric (Table 4; Figure 3). The recruitment of G_3_ individuals did not depend on population status or the origin of competitors (Table 5; Figure 4).

#### Relative survival and recruitment of both species in mixed cultures

The relative proportion of *P. acuta* in the G_2_ survivors at week 5, and in the G_3_ recruits at the end of the experiment, did not significantly depend on the allopatric versus sympatric nature of the pair of competing populations (X^2^_1df_ = 0.787 and 0.021 for survivors and recruits respectively, both P > 0.05), nor on the combination of statuses of the *A. marmorata* and *P. acuta* populations (X^2^_3df_ = 7.177 and 5.072, both P > 0.05). However, these results must be taken with caution given the low number of population pairs in each category. The proportion of *P. acuta* significantly exceeded ½ (intercept > 0 in logit scale in the binomial models) both in the G_2_ survivors (X^2^_1df_ = 6.241, P *=* 0.012) and in the G_3_ recruits (X^2^_1df_ = 20.702, P < 0.001) over all population pairs. The proportions fitted by the models were 0.62 +/-0.043 (SE) in G_2_ and 0.92 +/-0.035 in G_3_, corresponding to odd-ratios of 1.61 and 11.14 in favor of *P. acuta* for survival and recruitment respectively, and the corresponding raw means (+/-SE over experimental units) were 0.63 +/-0.038 and 0.80 +/-0.044 (odd-ratios of 1.72 and 4.10). Note that estimates and raw means differ because of the inclusion of random effects in the logit-scale binomial model. Frequencies of *P. acuta* in G_2_ at week 5 and in G_3_ recruits at week 16 were not significantly correlated across replicates (Spearman’s rho= 0.015, P > 0.05).

Summarizing the results, prediction (i_a) from Table 1 is fulfilled for growth (*P. acuta* competitors decrease growth in *A. marmorata* more than *A. marmorata* competitors, and the reverse is not true). For survival the prediction is not fulfilled: *P. acuta* seems insensitive to the identity of competitors, while naïve *A. marmorata* survive better with *P. acuta* than with an equal number of congeners. Prediction (i_b) is fulfilled, with a much higher recruitment of *P. acuta* in mixed cultures. Prediction (ii) is fulfilled for both survival (experienced *A marmorata* tolerate interspecific competition less well than naïve ones) and growth before age 2 weeks (idem); the tendency is similar but nonsignificant for size at 5 weeks. Regarding prediction (iii), the alternative (a) is fulfilled for growth (though not for survival) in both species, while none of the results are consistent with alternative (b).

## Discussion

We studied competitive interactions between the freshwater snails *A. marmorata* and *P. acuta* under laboratory conditions, and addressed three questions: (i) Is there a competitive asymmetry between the two species, and more specifically is the introduced species (*P. acuta*) competitively dominant over the resident (*A. marmorata*)? (ii) Does the resident species evolve to better resist competition, or rather to avoid it? (iii) Is there a population-specific component of coevolution between the two species at each site, such that sympatric populations of both species would tolerate each other better or worse than allopatric ones? We will address these three questions in turn.

### Asymmetrical competition, life-history syndromes and invasion

Our results first suggest that the resident species *A. marmorata* is less resistant to competition than the invasive one *P. acuta. Aplexa marmorata* individuals were indeed more affected by interspecific than by intraspecific competition, especially with regard to growth, and no such effect was detected in *P. acuta*. In addition, over the passage of generations (G_2_ and G_3_), the frequency of *A. marmorata* relative to *P. acuta* decreased in co-cultures; the former was even eliminated in some cases. This was due to a combination of higher adult survival and higher per-capita recruitment in *P. acuta*.

Competitive dominance reflects interspecific differences in traits that affect how they exploit resources and/or interfere with each other (Keddy, 2001). For instance, in plants, species growing faster to large size tend to monopolise light and outcompete slow-growing species (Hautier et al., 2009). We have previously shown that life-history traits differ between our two snail species in non-competitive laboratory conditions (Chapuis et al., 2017; a summary of these results is given in Appendix 1): compared to *A. marmorata, P. acuta* individuals reach sexual maturity slightly later, but are more fecund and survive much longer once adults, although their juvenile survival is lower. In freshwater snails, adults move faster and consume food more efficiently than juveniles (e.g., Selck et al., 2006). Thus, they efficiently inhibit the recruitment of juveniles in crowded conditions (pers. obs.), which is why we usually propagate snails by separating egg capsules from their parents. In our co-cultures, juvenile recruitment reflects the 5-week accumulation of many eggs with low individual probability of survival, probability that increased with time as adults progressively died out and were finally removed. Adults of *P. acuta* appear to monopolize resource, decreasing the growth of *A. marmorata* (hence, probably, their egg production); they also outlive *A. marmorata*, thus laying more eggs over time. Although *A. marmorata* eggs exhibit a higher hatching rate and survival in non-competitive conditions (Chapuis et al., 2017), this advantage seems insufficient to counterbalance the advantage of *P. acuta* during the adult phase. Our study thus corroborates the hypothesis proposed by Chapuis et al. (2017) that the life-history strategy of *P. acuta* provides an advantage in exploitative competition (a “K lifestyle”).

The conditions in our experiment (high density, single food source and unstructured aquarium environment) probably amplify the impacts of competition compared to natural situations. However, field data support the existence of asymmetrical effects of *P. acuta* on *A. marmorata* in Guadeloupe. First, population density in the latter decreased quickly in the Guadeloupe metacommunity after *P. acuta* arrival (Chapuis et al., 2017). Second, the presence of *P. acuta* in a site decreases its probability of being colonized by *A. marmorata* during the following year (Dubart et al., 2019; see summary of their results in Appendix 2). Reciprocally, the rate of local extinction of *P. acuta* is increased by the presence of *A. marmorata* in the site (Dubart et al., 2019; 2022). However, this rate remains lower than that of *A. marmorata*, resulting in a form of pre-emption at metacommunity / landscape scale: *P. acuta* colonizes new sites less often, but persists longer in each, and once established, tends to prevent recolonization by *A. marmorata*. As a consequence, stationary landscape occupancy by *A. marmorata* is predicted to decrease in Guadeloupe following the spread of *P. acuta*, though remaining far from the extinction threshold (Dubart et al., 2019).

Traits promoting competitive dominance therefore characterize the invasive *P. acuta*. The search for traits promoting biological invasions has been going on for a long time, with ambiguous results (e.g., Kueffer et al., 2013; Perkins & Nowak, 2013). This mixed outcome may be related to mixing different situations in which invasion is limited, or not, by competition (Krafts et al., 2007). Classical models of population growth (Elton 1946, Pimm 1991), have inspired the hypothesis that “fast” or “r-selected” lifestyles should be associated with invasion success, as suggested recently in amphibians, reptiles and mammals (Allen et al., 2017; Capellini et al., 2015). However, this may be the case only when rapid population growth is the only important limiting factor, i.e., in an “empty niche” situation. In the presence of a resident competitor, the limiting factor is to increase in frequency under competitive conditions. The associated trait syndrome should promote competitive dominance, at least relative to the resident species. For example, invasive birds tend to prioritize adult survival, and delay reproduction (review by Sol et al., 2012). Our snail example belongs to this category. Similar cases of asymmetric competition between related invasive and resident species (that may be themselves native or previously established invaders) have been observed in *Anolis* lizards (Culbertson et al., 2019) and fruit flies of the family Tephritidae (Duyck et al., 2006).

David et al. (2017) noticed that if negative species interactions, such as competition, act as important filters for potential invaders, those invasions that succeed are expected to have, on average, large negative impacts on resident species. As mentioned above, in the *P. acuta / A. marmorata* pair, such negative impacts are observed in the field (Dubart et al., 2019; 2022), though they seem relatively moderate compared to the outcome of laboratory experiments: *A. marmorata* does not seem to be at risk of extinction in Guadeloupe (while it does in our laboratory co-cultures). This situation may not be unusual, as extinctions due to competition by invasive species appear relatively rare compared to those resulting from predation or ecosystem engineering (Shine, 2010). Similarly, a recent report (IPBES, 2019) suggests that competition may play a somewhat lesser role than the combined effects of predation and herbivory. Instead of extinction, the outcome of invasive-resident competition is often niche partitioning. For example, resident Tephritid flies in the island of La Reunion were replaced by invaders in the most abundant resource (cultivated fruits in the lowland, e. g. mango) but persisted in marginal niches (peculiar families of fruits such as Solanacae, or highlands; Duyck et al., 2006; David et al., 2017; Facon et al., 2021). Similarly, in *Anolis* lizards, a native and an invasive species can co-exist owing to behavioral differences that promote differences in microhabitat use (Dufour et al., 2018). The generalist diet of freshwater snails (they basically graze on periphyton and exploit decayed organic matter) makes them little prone to specialization on different food sources – a form of specialization that has never been demonstrated in this group, though this would definitely deserve more research (see Dillon, 2000).

The kind of ‘niche partitioning’ that protects *A. marmorata* from extinction after invasion by *P. acuta* is more likely due to divergence in life-histories. The traits of *A. marmorata* (fast reproduction, high juvenile survival in the absence of competition, self-fertilization) characterize it as a “fugitive species” sensu Hutchinson (1951), as they may allow this species to exploit ephemeral resources (e.g., associated to macrophyte growth in ponds) before *P. acuta* numbers grow enough to outcompete it, which may eventually happen in places with stable resource levels. The natural environment of Guadeloupe is a system of patches (ponds) with frequent episodes of drought, eutrophication, or anoxia in the dry season, and rapid extinction-recolonization dynamics in freshwater snails (Pantel et al., 2022). In such a metapopulation context, the coexistence of the two species will depend on whether the disadvantage of *A. marmorata* in high-competition situations (e.g., adult populations limited by resource) is compensated by a better access to places where snail densities and competition are low, such as ponds refilled after a period of drought, or macrophyte stands regrowing after a perturbation. Indeed, metacommunity surveys show that *A. marmorata* more efficiently colonizes empty sites, but is more prone to local extinction, than *P. acuta* (Dubart et al., 2019; 2022; Appendix 2). The same study also shows that persistence of *A. marmorata* is facilitated, and colonization of *P. acuta* more difficult, when macrophyte cover is high (i.e., high resource levels). Thus, the different life-history syndromes of the two species are associated with demographic behaviors typical of, respectively, a better colonist (*A. marmorata*, the resident) and a better competitor (*P. acuta*, the invasive). The nature of the association is not fully clear however. The life-history trait differences measured here (Chapuis et al. 2017) probably directly contribute to competitive dominance by *P. acuta*, as this dominance is already evident in our simplified laboratory environment, with a single resource and no spatial structure (this study). Colonisation ability however was measured only in the field, and may depend not only on fecundity, growth and survival patterns (which may indeed favor fast population growth in *A. marmorata*), but also, crucially, on the ways by which snails disperse from one pond to the other. Little is known on this issue. Dispersal is passive in freshwater snails, and may occur either through aerial vectors such as birds, or through occasional water connexions between sites during floods (Van Leeuwen et al., 2013). Dubart et al. (2019) have shown that the invasive *P. acuta* is dependent on water connectivity to colonize new sites, while the resident *A. marmorata* efficiently colonizes both isolated and connected sites. However, it remains unclear which traits are responsible for this difference, and whether they belong to the set of traits measured in our experiment.

### Life history evolution and competition between resident and invasive species

Rapid evolution has been observed in several instances of interactions betwen invasive and resident species (Maron et al., 2004; Stuart et al., 2014; Chapuis et al., 2017). Competition can promote character displacement, i.e., trait evolution reducing impacts of competition between species, a process that occurs naturally during adaptive radiation (e.g., divergence in beak size among Darwin’s finches, Schluter & McPhail, 1992; Grant & Grant, 2006), as well as after invasions. For instance, in Dominica, the introduced lizard *Anolis cristatellus* and the native *Anolis oculatus* (Losos, 1990, Schluter & McPhail, 1993) diverged in body size after introduction, limiting niche overlap (review in Losos, 2009). However, laboratory environments offer highly simplified conditions and competition experiments usually do not reproduce the diversity of resources or niches that may promote character displacement in nature. This may lead to apparent contradictions between field and laboratory data. For example, in mosquitoes, different species were competitively dominant in the field (with litter as main food source) and in the laboratory (with artificial diet based on liver powder and yeast, Juliano, 1998). Similarly, reversed competitive dominance was observed in laboratory and field trials between thrips (*Frankliniella occidentalis* and *Thrips tabaci*), due to the differential susceptibilities of the species to insecticides present only in the field (Zhao et al., 2017). Thus, we might not necessarily expect that if the traits of our snail species had evolved to differenciate their niches and decrease competition in nature, they would also do so in our experimental conditions, with only one food source and a stable, unstructured environment.

Our previous study has shown that *A. marmorata* life-history traits have rapidly evolved in populations invaded by *P. acuta* (Chapuis et al., 2017): evolution has apparently favored traits associated with rapid population growth in non-competitive situations rather than exploitative competition, as their juvenile survival is enhanced while at the adult stage body size is smaller and life span shorter (Appendix 1). Our competition trials appear to validate this idea: the adult survival of experienced populations of *A. marmorata* is inferior to that of naive populations specifically when in competition with *P. acuta*. Thus *A. marmorata*, when reached by the *P. acuta* invasion, has evolved to become less competitive (at least in standard laboratory conditions). This apparent paradox, however, is solved under the hypothesis that the niche differenciation that allows *A. marmorata* and *P. acuta* to coexist is not a matter of resource type, but of ability to exploit bursts instead of low, stable levels of resource (see above). In invaded populations, the latter may be monopolized by *P. acuta*, and *A. marmorata* may have evolved to better exploit its residual niche (i.e., competition-free microhabitats). This situation could be considered as character displacement along the r-K gradient. Increased colonization ability may allow the resident species *A. marmorata* to bypass pre-emption of the resource by *P. acuta*. If *A. marmorata* reaches first empty ponds, it avoids being exposed to the inhibitory effect of established *P. acuta* detected in field surveys (Dubart et al. 2019, Appendix 2). More generally, our study illustrates the idea that interactions with invasive species may lead resident species to avoid rather than resist such interactions - another example is snakes that evolved a smaller gape to avoid predating poisonous invasive cane toads in Australia (Shine, 2010). Note, however, that similar recent studies in plants have found an evolution towards higher competitivity of the resident species (Germain et al., 2020) or an increase in allelopathy (Huang et al., 2018), suggesting that the direction of evolution (towards more or less competitive) may vary depending on the context. More studies are needed to assess whether these different outcomes of rapid evolution in resident species following invasion are generally related to differences in the initial degree of niche similarity and/or initial competitive asymmetry as expected from models (Godoy et al., 2014), or depends on other factors such as available genetic variation.

Invasive species may also evolve in response to interaction with residents, but our design is not able to detect this, as the “unevolved” state is not available in Guadeloupe. Indeed, all the *P. acuta* populations, be they recent (front) or ancient (core), have been founded by ancestors exposed to *A. marmorata* during the spread of *P. acuta* in Guadeloupe. Chapuis et al. (2017) did not detect important life-history differences between front and core populations – the small differences could be interpreted as possible impacts of temporary founder effects (invasion load; Colautti et al., 2006). In line with this finding, we only observed small differences between front and core populations of *P. acuta*, but no impact on competition: both population types were equally insensitive to the presence of *A. marmorata*, and outcompeted it in mixed cultures.

### Coevolution in sympatry

One of our most intriguing results is the stronger reduction in growth in both species by heterospecific competitors when the latter come from the same site (sympatry) than when they come from another invaded site (allopatry). This suggests that some specific evolution has taken place in one or both species within each site, which locally increases the impact of competition between the two. This coevolution pattern contrasts with the only other similar study, conducted in plants (Germain et al., 2020), as the ‘foreign’ versus ‘local’ nature of the competitors (allopatric versus sympatric pairs in our study) did not influence the outcome.

We must acknowledge that we do not have a straightforward explanation for this local coevolution. If each site harbours distinct environments, both species may have to evolve towards similar phenotypes to exploit it, and as a consequence of this similarity, their competitive impact on each other may increase in sympatric pairs (limiting similarity hypothesis). However, as soon as several niches or resource types are present within a given site, character displacement theory predicts that sympatric populations of resident and invasive species should diverge in resource use and minimize their reciprocal competitive impacts (Pfennig & Pfennig, 2009), rather than reinforce it. In addition, given that we provided a single food source, the evolution of resource exploitation efficiency cannot specifically affect the impact of the local population of competitors in our trials. Instead of exploitation, interference competition (Amarasekare, 2002; Huang et al., 2018) might be involved. This hypothesis is still speculative (pending further experiments), but Physids are known to alter their growth in response to scents of predators or congeners eaten by predators (Turner, 1997), such that they may also be able to chemically influence the growth of heterospecific competitors.

Whatever the mechanism, local co-evolution in our study system has increased competition impacts. This is in agreement with the hypothesis developed above, that the maintenance of the resident species in invaded ponds depends on its performances in occasional non-competitive situations rather than on standing interspecific competition in places where the invasive species is well established.

### Conclusion and perspectives

Species coexistence in metacommunity has been widely considered from an ecological perspective (Leibold & Chase, 2018), based on general frameworks such as that of Chesson (2000), highlighting either stabilizing mechanisms (based on trait differences that favor stable coexistence or reciprocal invasibility), or equalizing mechanisms (i.e., decreasing competitive asymmetry, up to neutrality) (Vanoverbeke et al., 2015; Leibold et al., 2019; Thompson et al., 2020). The role of evolutionary dynamics is usually not considered explicitly in this framework. Evolution can in principle reinforce both stabilizing and equalizing mechanisms; in competitive communities (i.e., within a guild) this will principally depend on whether it favors trait divergence or convergence among species.

Eco-evolutionary dynamics is not easy to detect empirically in stationary metacommunities, and is likely to proceed via small, population-specific trait changes (Leibold et al., 2019). In contrast, species invasions are large-scale perturbations that trigger large demographic and evolutionary adjustments, especially in resident species competing with newcomers (Haeuser et al., 2019). Under the coexistence perspective, successful invasion requires a “niche opportunity” (Shea & Chesson, 2002; Pearson et al. 2018; Latombe et al. 2021), representing the invasibility of the community by a new set of traits. In guilds of species with similar resource ranges, we argued that this condition often selects invasive species whose traits place them above residents in the competitive hierarchy (David et al., 2017). Our study provides an example where the competition between invasive and resident species is clearly asymmetrical, in favor of the former, as expected from the phylogenetic and trophic similarity between the two, and the hypothesis of a competition filter to invasion. Contrary to other examples (Huang et al., 2018; Germain et al., 2020), rapid evolution in the resident species has tended to reinforce the initial competitive asymmetry. This is consistent with a mode of coexistence based on a form of niche partitioning along the colonization-competition (‘r-K’) axis, whereby the resident species specializes in exploiting ephemeral low-competition situations, being otherwise outcompeted by the invasive species. This is akin to cases of character displacement in other animals (Feiner et al., 2021; Grant & Grant, 2006), except that the displaced characters are life-history traits rather than traits related to the exploitation of different resource types (Chapuis et al., 2017). As in most published examples of invasive-resident interactions, we have no reason to believe that evolution was absolutely necessary for the resident to persist in the presence of its new competitor (i.e., evolutionary rescue sensu stricto; Carlson et al., 2014); but several other examples suggest that rapid evolution of competitive interactions is commonplace after invasions (Huang et al., 2018; Germain et al., 2020). Coming back to Chesson’s framework, in our system, evolution has tended to reinforce stabilizing mechanisms (a difference in trait that allows the competitively inferior resident species to survive competition by the invader, at the metapopulation scale), thus making their regional coexistence even more secure than it initially was. This is not necessarily the only possible outcome; evolution might, in other contexts, decrease competitive asymmetry between invasive and resident species, i. e. have an equalizing rather than stabilizing effect. Understanding under which conditions evolution tends to equalize fitnesses by increasing the resistance to competition (such as in plants: Huang et al., 2018; Germain et al., 2020), rather than promoting its avoidance (as in snails) is a worthwile goal for future research.

## Appendix 1

Mean values (sd) of several phenotypic traits studied in non-competitive laboratory conditions in Chapuis et al. (2017). SM = sexual maturity. The biovolume was estimated based on an ellipsoid considering shell length and width meaured as estimates of its height and diameter respectively, and fecundity was estimated over two weeks.

|  | Age at SM<br>(days) | Body length<br>at SM (mm) | Biovolume at<br>SM (mm<sup>3</sup>) | Fecundity<br>(egg number) | Age at death<br>(days) | Juvenile<br>survival (%) |
| --- | --- | --- | --- | --- | --- | --- |
| *Aplexa marmorata* | 28.14<br>(4.82) | 6.70<br>(0.84) | 39.74<br>(15.12) | 135.63 (40.72) | 52.60 (26.83) | 74.92<br>(24.40) |
| *Physa acuta* | 30.77<br>(3.18) | 6.23<br>(0.65) | 40.91<br>(13.15) | 171.64 (72.17) | 78.30 (22.96) | 52.46<br>(26.85) |

## Appendix 2

Colonisation and extinction rates in ponds in Guadeloupe, for the resident species *A. marmorata*, and the invasive species *P. acuta*, in the presence and absence of the other species. These rates have been estimated by Dubart et al. (2019) based on a 17-year survey of presence-absence of the two species in 250 sites every year, some of them with replicate visits within a season. Estimates are from state-space models accounting for imperfect detection (i.e., fitting a fraction of sites where the species is present but not detected). The estimates apply to ponds with water, and average environmental characteristics; the species also persist in the soil in dessicated ponds (the model does not assume they are extinct when the pond is found dry). *p*_*c*_ and *p*_*e*_ are estimated probabilities that a site gets colonised or extinct in one year, with associated 95% credibility interval.

| | $p_c$ (other species<br>absent) | $p_c$ (other species<br>present) | $p_e$ (other species<br>absent) | $p_e$ (other species<br>present) |
| --- | --- | --- | --- | --- |
| *A. marmorata* | 0.71<br>(0.61-0.82) | 0.325<br>(0.26-0.40) | 0.25<br>(0.125-0.48) | 0.25<br>(0.18-0.345) |
| *P. acuta* | 0.51<br>(0.33-0.68) | 0.37<br>(0.295-0.44) | 0.01<br>(0.00-0.055) | 0.15<br>(0.045-0.26) |

## Acknowledgements

We thank Thomas Lamy and Jean-Pierre Pointier for their help with field sampling. And the late Violette Sarda for help with husbandry maintenance. We also thank recommender B. Phillips, D. Reznick and two anonymous reviewers for their useful comments.

## Data, scripts, code, and supplementary information availability

Data are available online: <u>10.5281/zenodo.10041279</u>.

Scripts and code are available online: <u>10.5281/zenodo.10041344</u>.

